# Deep-sea corals near cold seeps associate with chemoautotrophic bacteria that are related to the symbionts of cold seep and hydrothermal vent mussels

**DOI:** 10.1101/2020.02.27.968453

**Authors:** Samuel A. Vohsen, Harald R. Gruber-Vodicka, Eslam O. Osman, Matthew A. Saxton, Samantha B. Joye, Nicole Dubilier, Charles R. Fisher, Iliana B. Baums

**Affiliations:** The Pennsylvania State University; Max Planck Institute for Marine Microbiology; University of Georgia

**Keywords:** Deep-sea corals, SUP05, chemoautotrophy, seeps, sulfur-oxidizer

## Abstract

Cnidarians are known for their symbiotic relationships, yet no known association exists between corals and chemoautotrophic microbes. Deep-sea corals, which support diverse animal communities in the Gulf of Mexico, are often found on authigenic carbonate in association with cold seeps. Sulfur-oxidizing chemoautotrophic bacteria of the SUP05 cluster are dominant symbionts of bathymodiolin mussels at cold seeps and hydrothermal vents around the world and have also been found in association with sponges. Therefore, we investigated whether other basal metazoans, corals, also associate with bacteria of the SUP05 cluster and report here that such associations are widespread. This was unexpected because it has been proposed that cnidarians would not form symbioses with chemoautotrophic bacteria due to their high oxygen demand and their lack of specialized respiratory structures. We screened corals, water, and sediment for SUP05 using 16S metabarcoding and found SUP05 phylotypes associated with corals at high relative abundance (10 – 91%). These coral-associated SUP05 phylotypes were coral host specific, absent in water samples, and rare or not detected in sediment samples. The genome of one SUP05 phylotype associated with *Paramuricea* sp. type B3, contained the genetic potential to oxidize reduced sulfur compounds and fix carbon and these pathways were transcriptionally active. Finally, the relative abundance of this SUP05 phylotype was positively correlated with chemoautotrophically-derived carbon and nitrogen input into the coral holobiont based on stable carbon and nitrogen isotopic compositions. We propose that SUP05 may supplement the diet of its host coral through chemoautotrophy or may provide nitrogen, essential amino acids, or vitamins. This is the first documented association between a chemoautotrophic symbiont and a cnidarian, broadening the known symbioses of corals and may represent a novel interaction between coral communities and cold seeps.

## Introduction

Deep-sea corals are important foundation species that support a diverse community of animals that rivals the diversity of shallow, tropical reefs and includes many commercially important fish species (Costello et al. 2005; Buhl-Mortensen and Mortensen 2005; Jensen and Frederiksen 1992; Cordes et al. 2008; Lessard-Pilon et al. 2010). Deep-sea coral communities can be found along all continental margins from the Arctic to the Antarctic and on seamounts worldwide (Yesson et al. 2012; Roberts et al. 2006). In the deep Gulf of Mexico, the ranges of many deep-sea coral species overlap with cold seeps characterized by elevated concentrations of hydrogen sulfide and/or hydrocarbons ranging from methane to crude oil (Quattrini et al. 2013; Becker et al. 2009; Cordes et al. 2008).

Some animals at cold seeps form associations with chemoautotrophic symbionts which provide them with nutrition from the oxidation of these reduced chemical species (Fisher et al. 1993; Childress et al. 1986; Brooks et al. 1987). Bathymodiolin mussels and vestimentiferan tubeworms obtain the bulk of their nutrition from these symbionts and form dense assemblages which in turn support highly productive animal communities (Bergquist et al. 2003; Bergquist et al. 2005; Sibuet and Olu 1998). The sulfur-oxidizing symbionts of mussels that make these communities possible belong to the widespread SUP05 cluster of gamma-proteobacteria (Petersen et al. 2012). This cluster includes sulfur-oxidizing symbionts found in both cold seep and hydrothermal vent fauna (Petersen et al. 2012). The SUP05 cluster also includes many free-living species which are abundant at oxygen minimum zones and hydrothermal vents where they dominate dark carbon fixation in these major carbon sinks (Mattes et al. 2013; Shah et al. 2019; Walsh et al. 2009; Anantharaman et al. 2013; Glaubitz et al. 2013).

Early work with the mound-forming deep-sea scleractinian coral, *Lophelia pertusa*, found that coral mounds formed near cold seeps where methane concentrations in the sediment were elevated. Initial hypotheses included that corals fed on chemoautotrophic and methanotrophic bacteria from the seeps (Hovland and Thomsen 1997; Hovland and Risk 2003). Analyses of the microbial community associated with *Lophelia pertusa* identified a relative of the SUP05 cluster that was present in corals from Norway and the Gulf of Mexico (Kellogg et al. 2009; Neulinger et al. 2008). However, microscopic analysis did not locate these bacteria and showed no evidence of an abundant bacterial associate in corals (Neulinger et al. 2009). Other work analyzing available particulate matter and stable isotopes demonstrated that *Lophelia pertusa* does not receive detectable nutritional input from seep activity and simply use authigenic carbonates produced at cold seeps as substrate after seepage has waned (Becker et al. 2009; Kiriakoulakis et al. 2004). These results and the oxygen demand of chemoautotrophic symbionts has led some to suggest that cnidarians would not form associations with sulfide-oxidizing symbionts since they have no specialized respiratory structures or oxygen transport mechanisms (Childress and Girguis 2011).

Here we report the discovery of an association between deep-sea corals and members of the SUP05 cluster. We used 16S metabarcoding to screen corals, water, and sediment for the presence of members of the SUP05 cluster. We screened 421 colonies from 42 coral morphospecies including scleractinians, octocorals, antipatharians, and zoanthids from deep-sea sites and included mesophotic and shallow-water coral species for comparison. We then focused on one species, *Paramuricea* sp. type B3 (Doughty et al. 2014), from one site with seepage, BOEM lease block AT357, as a model to study the role of SUP05 in corals. We sequenced the metagenome and metatranscriptome of *Paramuricea* sp. type B3 to assemble the genome of its associated SUP05 and to investigate its metabolic potential and to determine which genes and pathways were expressed. Finally, we investigated the nutritional relationship between *Paramuricea* sp. type B3 and its SUP05 using carbon and nitrogen stable isotopic analysis.

## Methods

### Collections

Four hundred and twenty-one coral colonies belonging to forty-two morphospecies were collected from thirty-one deep-sea and mesophotic sites in the Gulf of Mexico during eight cruises from 2009 to 2017 (Table 1, Fig. 1). Signs of seepage were restricted to depths over 400 m and sites were heterogeneous with respect to signs of active seepage.

**Table 1.**
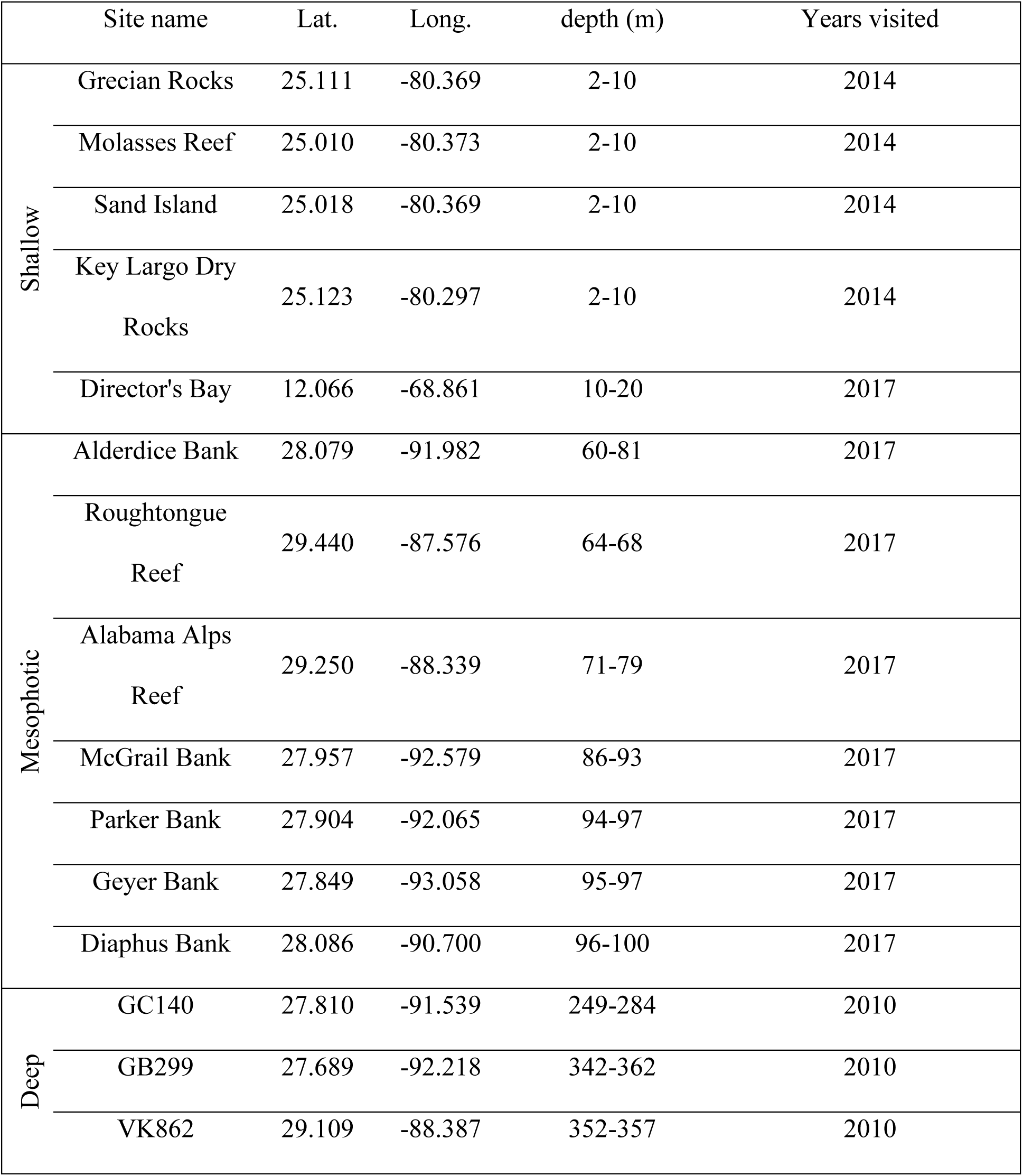
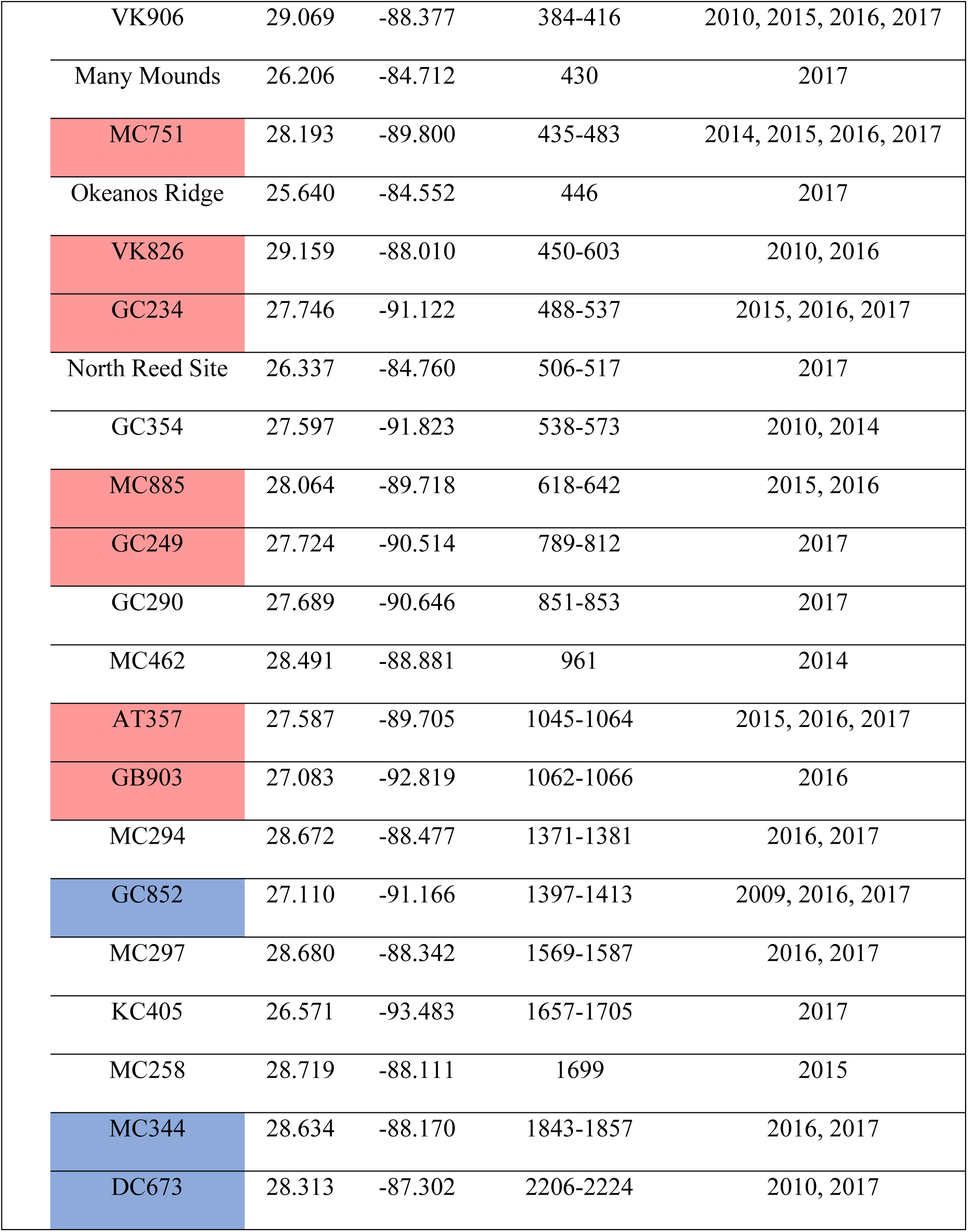
Sampling locations and years sampled. Sites where corals were sampled near signs of seepage are highlighted in salmon. Sites within lease blocks with active seepage but not observed near sampled corals are highlighted in blue.

|  | Site name | Lat. | Long. | depth (m) | Years visited |
| --- | --- | --- | --- | --- | --- |
| Shallow | Grecian Rocks | 25.111 | -80.369 | 2-10 | 2014 |
|  | Molasses Reef | 25.010 | -80.373 | 2-10 | 2014 |
|  | Sand Island | 25.018 | -80.369 | 2-10 | 2014 |
|  | Key Largo Dry<br>Rocks | 25.123 | -80.297 | 2-10 | 2014 |
|  | Director's Bay | 12.066 | -68.861 | 10-20 | 2017 |
| Mesophotic | Alderdice Bank | 28.079 | -91.982 | 60-81 | 2017 |
|  | Roughtongue<br>Reef | 29.440 | -87.576 | 64-68 | 2017 |
|  | Alabama Alps<br>Reef | 29.250 | -88.339 | 71-79 | 2017 |
|  | McGrail Bank | 27.957 | -92.579 | 86-93 | 2017 |
|  | Parker Bank | 27.904 | -92.065 | 94-97 | 2017 |
|  | Geyer Bank | 27.849 | -93.058 | 95-97 | 2017 |
|  | Diaphus Bank | 28.086 | -90.700 | 96-100 | 2017 |
| Deep | GC140 | 27.810 | -91.539 | 249-284 | 2010 |
|  | GB299 | 27.689 | -92.218 | 342-362 | 2010 |
|  | VK862 | 29.109 | -88.387 | 352-357 | 2010 |

|  |  |  |  |  |
| --- | --- | --- | --- | --- |
| VK906 | 29.069 | -88.377 | 384-416 | 2010, 2015, 2016, 2017 |
| Many Mounds | 26.206 | -84.712 | 430 | 2017 |
| MC751 | 28.193 | -89.800 | 435-483 | 2014, 2015, 2016, 2017 |
| Okeanos Ridge | 25.640 | -84.552 | 446 | 2017 |
| VK826 | 29.159 | -88.010 | 450-603 | 2010, 2016 |
| GC234 | 27.746 | -91.122 | 488-537 | 2015, 2016, 2017 |
| North Reed Site | 26.337 | -84.760 | 506-517 | 2017 |
| GC354 | 27.597 | -91.823 | 538-573 | 2010, 2014 |
| MC885 | 28.064 | -89.718 | 618-642 | 2015, 2016 |
| GC249 | 27.724 | -90.514 | 789-812 | 2017 |
| GC290 | 27.689 | -90.646 | 851-853 | 2017 |
| MC462 | 28.491 | -88.881 | 961 | 2014 |
| AT357 | 27.587 | -89.705 | 1045-1064 | 2015, 2016, 2017 |
| GB903 | 27.083 | -92.819 | 1062-1066 | 2016 |
| MC294 | 28.672 | -88.477 | 1371-1381 | 2016, 2017 |
| GC852 | 27.110 | -91.166 | 1397-1413 | 2009, 2016, 2017 |
| MC297 | 28.680 | -88.342 | 1569-1587 | 2016, 2017 |
| KC405 | 26.571 | -93.483 | 1657-1705 | 2017 |
| MC258 | 28.719 | -88.111 | 1699 | 2015 |
| MC344 | 28.634 | -88.170 | 1843-1857 | 2016, 2017 |
| DC673 | 28.313 | -87.302 | 2206-2224 | 2010, 2017 |

**Figure 1.**
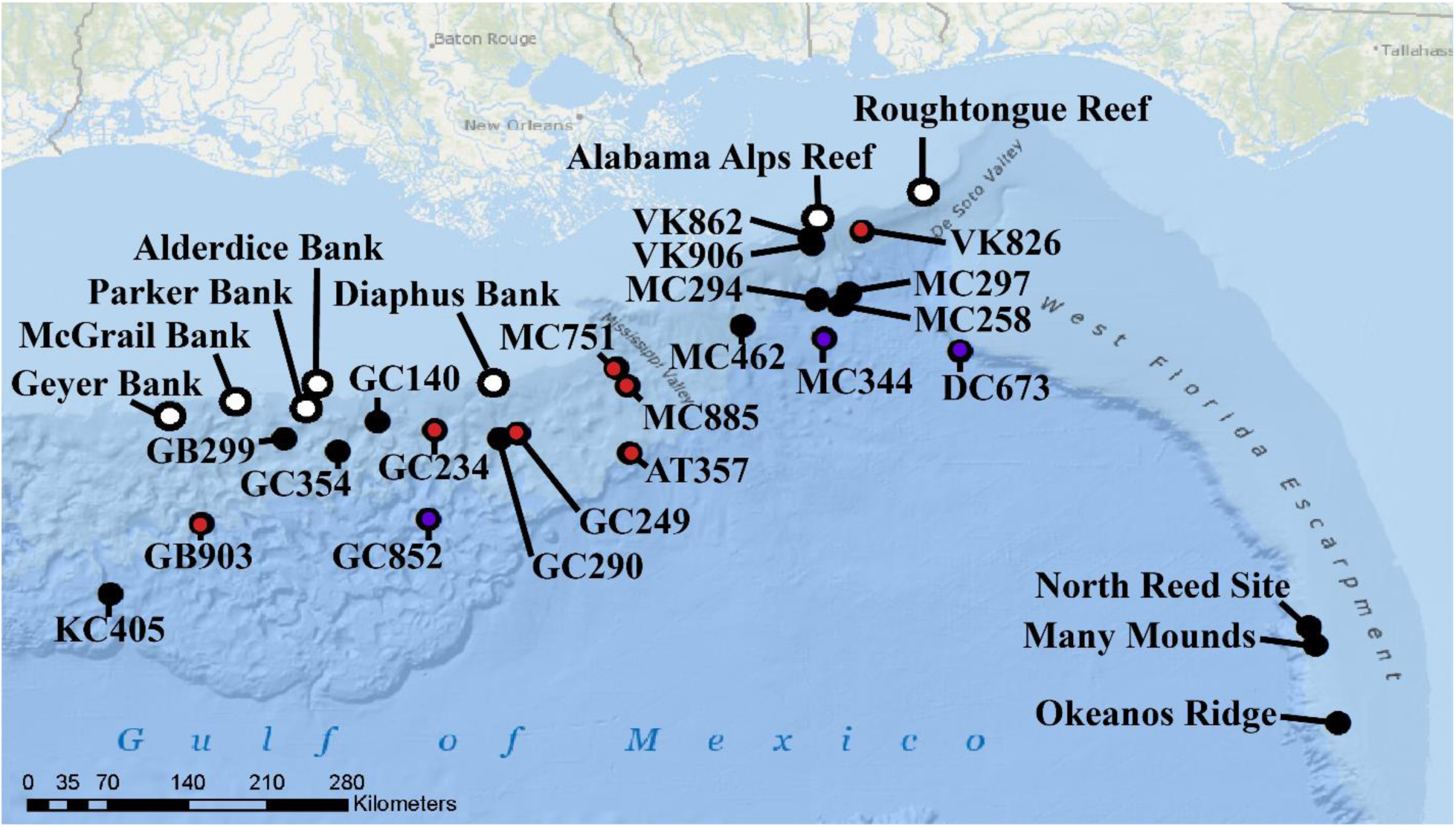
Map of sampling locations. Deep-sea sites (>200 m) are designated with black dots. Mesophotic sites (50 – 100m) are designated with white dots. Sites where corals were sampled near signs of seepage are marked with a red dot. Sites with known signs of seepage within the lease block but not observed near corals are marked with a blue dot. Bathymetric map obtained from BOEM (credit: William Shedd, Kody Kramer)

Deep-sea corals were sampled using specially designed coral cutters mounted on the manipulator arm of remotely operated vehicles (ROVs). Coral fragments were removed, placed in temperature insulated containers until recovery of the ROV, and were maintained at 4 °C for up to 4 hours until preservation in ethanol or frozen in liquid nitrogen. Corals were also sampled from 4 shallow-water sites in the Florida Keys and 1 in Curaçao from 1 – 20 m depth. Coral fragments were removed using either a hammer and chisel or bonecutters and placed in separate Ziploc bags. The shallow water coral samples were preserved in ethanol or frozen in liquid nitrogen on the boat or on shore within 8 hours.

Sediment samples were taken using an ROV in close proximity to many of the deep-sea coral collections with 6.3 cm diameter push cores. Upon recovery of the ROV, 1 mL of sediment from the top 1 cm of each sediment core was frozen in liquid nitrogen for microbiome analysis. In 2015, water was sampled using a Large Volume Water Transfer System (McLane Laboratories Inc., Falmouth, MA) which filtered approximately 400 L of water through a 0.22 micron porosity filter (142 mm diameter). One quarter of each filter was preserved in ethanol and another quarter was frozen in liquid nitrogen for these analyses. In 2016 and 2017, 2.5 L niskin bottles were fired above corals and upon recovery were filtered through 0.22 micron filters which were frozen in liquid nitrogen.

### Description of study site: AT357

A large community of corals, dominated by *Paramuricea* sp. type B3 and *Madrepora oculata* is present in BOEM lease block AT357 (27.587 N, −89.705 W) between 1045 – 1064 m depth (Fig. 2). Corals from this site were used to investigate potential nutritional interactions between corals and their SUPO5 symbionts. Corals are found over an area of at least 250 by 250 m at this site and there are numerous areas of active seepage interspersed among the coral colonies (Fig. 2). Colonies of *Paramuricea* sp. type B3 were found on mounds of *Madrepora oculata*, and on authigenic carbonates formed in areas of previous or current active seepage. Large bacterial mats were often present in small troughs and depressions between the corals. Chemoautotrophic *Bathymodiolus brooksi* mussels and vesicomyid clams were occasionally present in these areas of active seepage as well. Twenty-two *Paramuricea* sp. type B3 colonies were sampled across this site including colonies growing near active seeps and others from areas with no visible indications of currently active seepage within 10 m.

**Figure 2.**
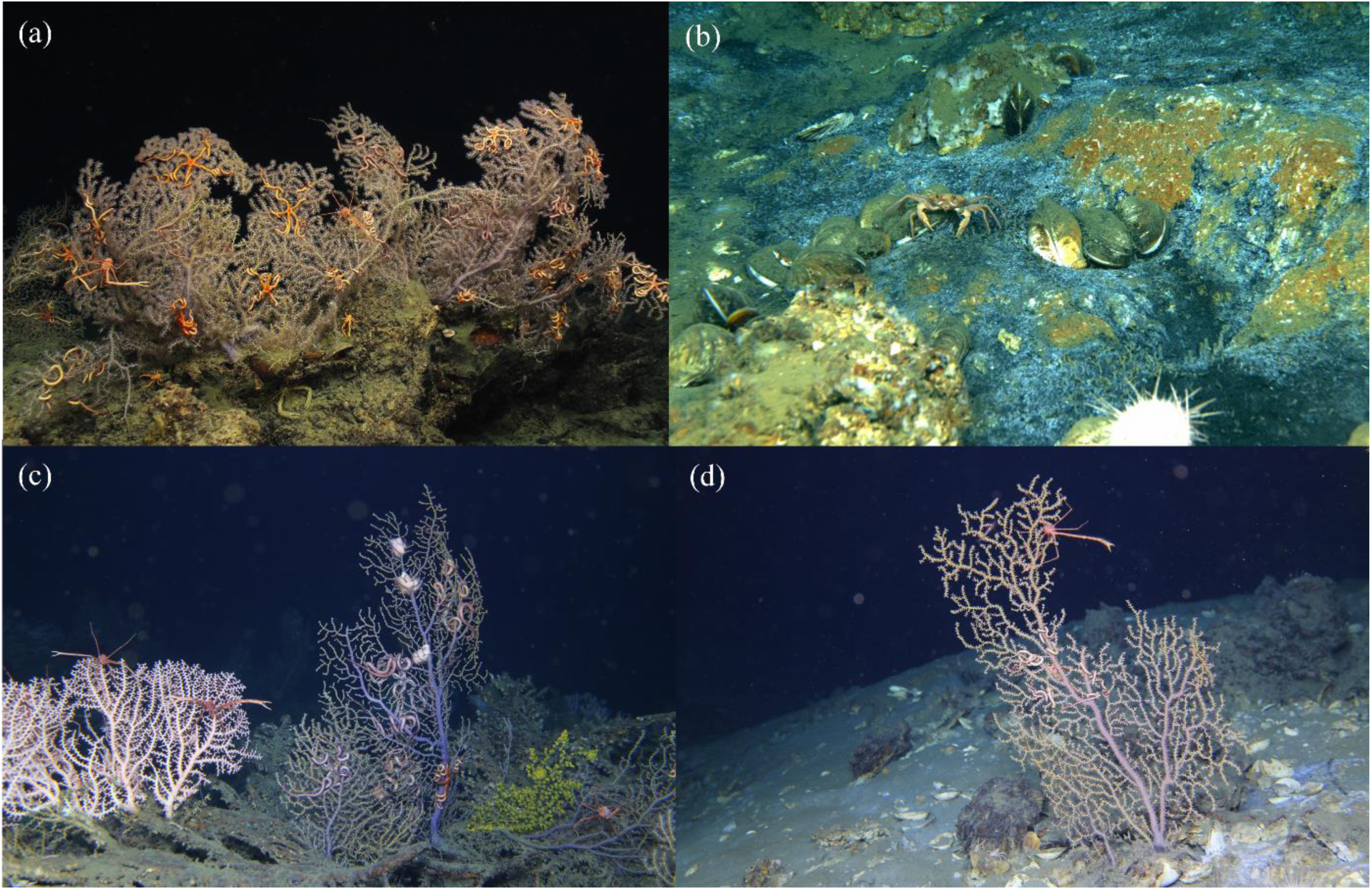
Images at AT357 of *Paramuricea* sp. type B3 and associated fauna (a), bacterial mats and living chemosymbiotic mussels (*Bathymodiolus brooksi*) indicating active seepage of reduced sulfur species (b), *Madrepora oculata* and zoanthid gold corals co-occurring with *Paramuricea* sp. type B3 (c), and colonies of *Paramuricea* sp. type B3 growing very near bacterial mats (d).

### 16S metabarcoding

Corals, water, and sediment were screened for SUP05 using 16S metabarcoding. DNA was extracted from coral tissue and sediment samples using DNeasy PowerSoil kits (Qiagen, Hilden, Germany) following manufacturers protocols using approximately 1 cm of coral branches and about 0.25 g of sediment. DNeasy PowerSoil kits were also used for water samples collected in 2015 using 1 cm^2^ of filter. Replicate extractions were performed on quarters of the filter preserved in both ethanol and frozen in liquid nitrogen for all water samples in 2015. DNA was extracted from all other water samples using Qiagen DNeasy PowerWater kits.

The V1 and V2 region of the 16S rRNA gene were amplified using universal bacterial primers 27F and 355R (Rodriguez-Lanetty et al. 2013) with Fluidigm CS1 and CS2 adapters (Illumina, San Diego, CA). PCR was conducted with the following reaction conditions: 0.1 U/µL Gotaq (Promega, Madison, WI), 1X Gotaq buffer, 0.25mM of each dNTP, 2.5 mM MgCl_2_, and 0.25 µM of each primer, and the following thermocycler conditions: 95 °C for 5 min; 30 cycles of 95 °C for 30 sec, 51 °C for 1 min, and 72 °C for 1 min; and finally 72 °C for 7 min. Amplification was confirmed on a 1% LE agarose gel. Libraries were prepared by the University of Illinois Chicago DNA services facility and sequenced on two separate runs on an Illumina Miseq platform.

Amplicon sequence data were analyzed using QIIME2 (ver2017.11). Reads were joined using vsearch and quality filtered using q-score-joined. Deblur denoise was used to detect chimeras, construct sub-operational taxonomic units (later referred to as phylotypes), and trim assembled reads to 300bp. Classifications of phylotypes was achieved by building a classifier with the SILVA v128 SSU 99% consensus sequence database using a naïve Bayes fit. Phylotypes classified as SUP05 cluster were used for later analyses.

### 16S rRNA cloning for SUP05

Cloning was employed to obtain the full length 16S rRNA gene of SUP05 phylotypes in coral samples with the highest relative abundance of SUP05 in their microbiome. To amplify the full length bacterial 16S rRNA gene, we used the universal bacterial primers 27F and 1492F with the following PCR conditions on a Mastercycler pro (Eppendorf, Hamburg, Germany); 95 °C for 5 min for DNA denaturation; 30 cycles of 95 °C for 1 min, 55 °C for 1 min sec and 72 °C for 1 min; followed by a final extension at 72 °C for 10 min. PCR reactions consisted of 1X Gotaq buffer (Promega, Madison, WI), 2.5 mM MgCl_2_, 0.25 mM dNTPs, 0.4 µM of each primer, and 0.1 U/µL Gotaq (Promega) and 1 or 4 µL DNA extract in 25 µL reaction volume. PCR amplicon size was visualized on a 1% LE agarose gel (110 V for 35 min) against a size standard to confirm amplification of the correct target sequence. The PCR products were cleaned using ExoSAP-IT Express cleanup kits (Thermo Fisher, Waltham, MA) and used as templates in a TOPO TA cloning reaction (Thermo Fisher, Waltham, MA). The TOPO cloning reaction combined 4 µl of cleaned PCR product, 1µl of salt buffer, and 1µl of TOPO TA vector, and was incubated at room temperature for 10 minutes. Transformation was conducted using One Shot™ TOP10 Chemically Competent *E. coli* (Invitrogen, USA) by combining 3µl of TOPO cloning reaction with 50 µL of *E. coli* cells. Samples were placed on ice for 30 min, heat shocked in a water bath at 42 °C for 30 sec, and finally placed back on ice for 2 min. Transformed cells were mixed with 250 µL of S.O.C. medium and incubated in 37 °C for 1 hour with shaking (220 rpm). After incubation, 50 - 150 µL of cultured cells were plated onto LB agar medium containing 50 µL X-GAL (40 mg/mL) and 50 mg kanamycin per plate and incubated for 15 h at 37 °C. Later, 8-10 white colonies were each placed in 20 µl sterilized water and 1 µl was taken as a template for final PCR amplification with either i) 27F and 1492R primers using the same PCR conditions as above except an increased initial denaturation of 10 min or ii) M13 forward and reverse primers using an initial denaturation at 95 °C for 10 min; 32 cycles of 95 °C for 30 s, 55 °C for 30 s, and 72 °C for 45 s; and final extension at 72 °C for 10 min. Correct amplicon size was confirmed on a 1% agarose gel. PCR products were cleaned with ExoSAP-IT Express cleanup kits and sent to the genomics core facility at Pennsylvania State University for Sanger sequencing with either 27F/1492R or M13 primer sets.

### Phylogenetic Analyses

A phylogenetic tree was constructed to infer the evolutionary positions of the coral-associated SUP05 phylotypes using both partial and full length 16S rRNA gene sequences produced in this study combined with sequences from public databases. Sequences were aligned using MUSCLE ver3.8.425 (Edgar 2004) and a maximum likelihood tree was constructed using iqtree ver1.6.10 (Nguyen et al. 2014). ModelFinder (Kalyaanamoorthy et al. 2017) was used to choose a transversion model with equal base frequencies and a FreeRate model for rate heterogeneity across sites with 3 categories (TVMe+R3) based on the highest BIC score. UFBoot2 (Hoang et al. 2017) was used to obtain ultrafast bootstrap support values (UFBoot) using 1000 replicates.

### Stable Isotope Analysis

Tissue samples from twenty-two colonies of *Paramuricea* sp. type B3 from AT357 were processed for stable isotopic analysis to investigate any potential trophic relationship between the coral host and their associated SUP05. Coral samples were collected in 2015 and 2016, frozen in liquid nitrogen upon recovery of the ROV, and stored at −80 °C for up to 12 months. Frozen tissue was dried at 47 °C for two days then acidified with 2-5 drops of 2 N phosphoric acid to remove calcium carbonate. Samples were then redried and acidified until all carbonate was removed as indicated by a lack of bubbling and then redried a final time. Two mg from each dried sample was encapsulated in tin and sent to UC Davis for carbon and nitrogen stable isotopic analysis. Samples were analyzed using a PDZ Europa ANCA-GSL elemental analyzer coupled to a PDZ Europa 20-20 isotope ratio mass spectrometer (Sercon, Cheshire, UK).

A linear model was constructed to test the correlation between the relative abundance of SUP05 in *Paramuricea* sp. type B3 and the stable isotopic composition of the coral tissue using sampling year as a categorical variable.

### Metagenomes and Metatranscriptomes

The genome and transcriptome of a coral-associated member of the SUP05 cluster were sequenced in order to assess its metabolic potential.

DNA was extracted from four *Paramuricea* sp. type B3 colonies using DNA/RNA allprep kits (Qiagen, Hilden, Germany) with β-mercaptoethanol added to the RLT+ lysis buffer following the manufacturer’s instructions. Additionally, RNA was extracted from two of those *Paramuricea* sp. type B3 colonies using an RNeasy extraction kit (Qiagen, Hilden, Germany). The RNA extracts were enriched for bacteria by depleting coral host rRNA using a Ribo-Zero Gold Yeast kit (Illumina, San Diego, CA, USA) following the manufacturer’s protocols.

DNA extracts were sent to the Max Planck Genome Center in Cologne, Germany for library preparation and were sequenced on a HISEQ2500 platform using 150 bp paired-end reads. The two *Paramuricea* sp. type B3 libraries with the highest coverage of the SUP05 genome were sequenced a second time for higher coverage of the SUP05 genome. The RNA extracts were submitted to the Penn State Genomics Core for library preparation and were sequenced on a MISEQ platform using 100 bp single-end reads.

All DNA sequence libraries were screened for SUP05 SSU rRNA using phyloFlash ver3.3 (Gruber-Vodicka et al. 2019). The library with the highest coverage of SUP05 SSU rRNA was assembled using megahit ver1.1.2 (Li et al. 2015). The reads from each of the four libraries were individually mapped to this assembly using bbmap with default parameters and these coverages were used to bin the assembly using metabat (ver2.12.1). A bin associated with SUP05 and all contigs connected in the assembly were used as a draft genome and were annotated using RASTtk ver2.0 (Aziz et al. 2008; Overbeek et al. 2005). The two RNA sequence libraries were mapped to these gene annotations using kallisto ver0.44.0 (Bray et al. 2016) to determine which genes were expressed using 100 bootstrap replicates, an estimated fragment length of 180 with a standard deviation of 20, and an index with a k-mer size of 31.

## Results

### Occurrence of SUP05 cluster in corals, water, and sediment

A total of 72 SUP05 phylotypes were identified throughout the whole dataset and were present in coral, water, and sediment samples. We considered a phylotype to be abundant in a coral if it constituted >10% of the microbial community in at least one coral sample. Fifteen SUP05 phylotypes were abundant in coral samples ranging between 12 and 91% of the microbial community (Fig. 3, Table S1). These phylotypes were abundant in 13 coral morphospecies from a wide depth range (∼350 - 1800 m) including (ordered by depth of occurrence) *Paramuricea* sp., *Muriceides hirta, Paragorgia* sp. 2, *Swiftia* sp., *Acanthogorgia aspera, Callogorgia delta*, unidentified bamboo 1, *Chrysogorgia* sp., *Paramuricea* sp. type B3, *Corallium* sp., *Swiftia pallida, Paramuricea biscaya*, and *Stichopathes* sp. 3 from sites in BOEM lease blocks VK906, MC751, VK826, GC234, MC462, AT357, GB903, GC852, and MC344 (Fig. 3).

**Figure 3.**
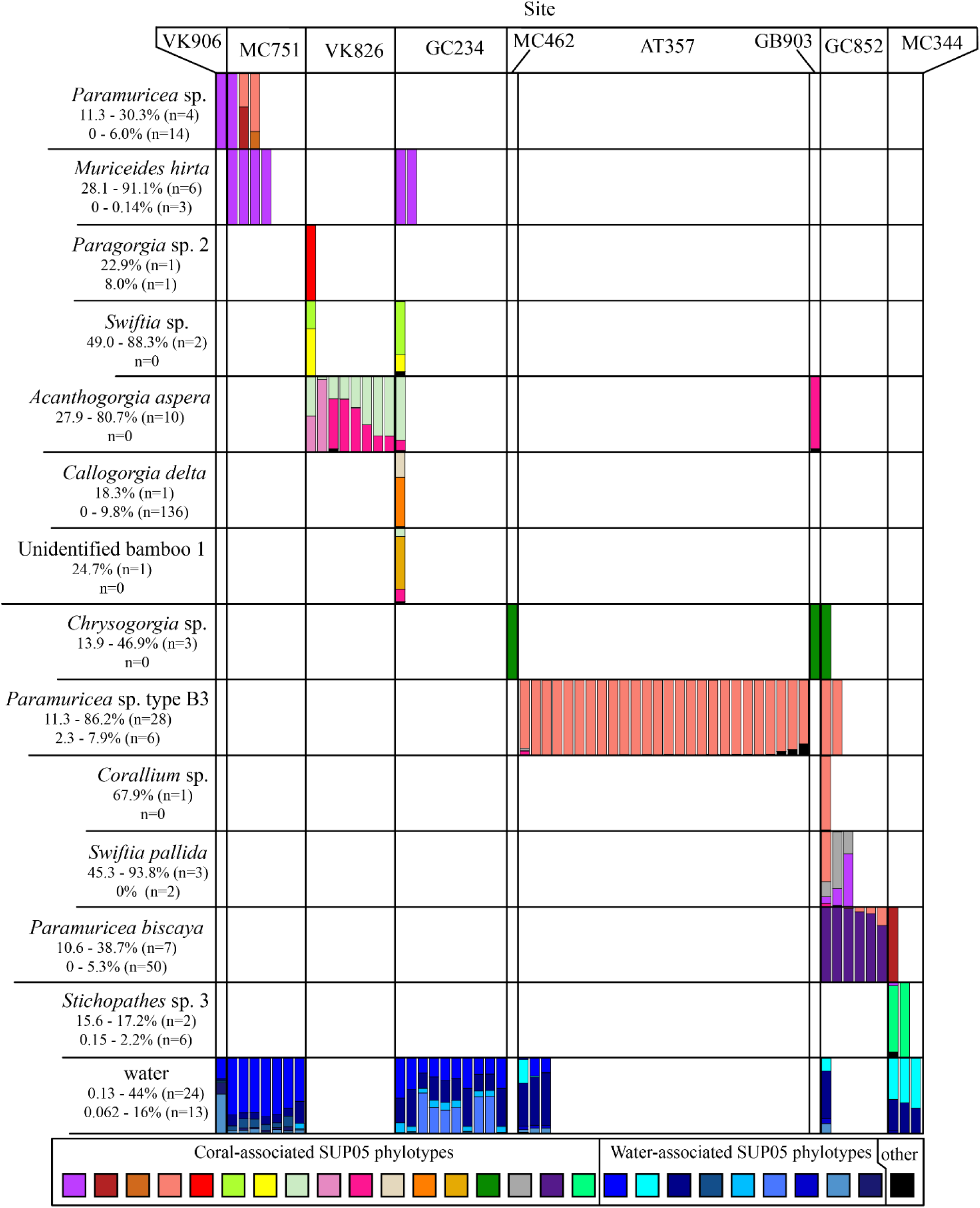
Occurrence of SUP05 phylotypes in coral species with higher than 10% SUP05. All water samples from sites with corals that met this criterion were included. Each bar represents the composition of SUP05 phylotypes in an individual coral or water sample. SUP05 phylotypes that comprise less than 10% and don’t occur in water samples were pooled and all colored black. Samples from the same morphospecies are organized in rows and those from the same site are organized into columns. Reported below each morphospecies name is the range of relative abundances of all SUP05 phylotypes for each sample shown in the figure followed by the ranges for all samples not shown.

These abundant, coral-associated SUP05 phylotypes exhibited specificity for coral hosts. Twelve phylotypes were only abundant in corals within a single genus or species and were absent or rare in other corals (Fig. 3). Further, many coral species retained the same dominant SUP05 phylotype across multiple sites while other coral species at the same sites hosted different SUP05 phylotypes. For example, the five co-occurring corals at site GC234 each hosted different dominant SUP05 phylotypes.

None of these abundant coral-associated SUP05 phylotypes were ever detected in a water sample, and nine of the twelve were not detected in any sediment sample. Those present in sediment samples comprised a maximum of less than 0.2% of the sediment microbial community (Table S1). In addition, 58 other SUP05 phylotypes were detected that comprised less than 10% of the microbial community in individual corals. Of these, 29 were found only in coral samples and never in water or sediment samples (Table S1). These phylotypes were associated with *Acanthogorgia aspera, Bathypathes* sp. 1, *Callogorgia americana, Callogorgia delta, Chelidonisis aurantiaca, Clavularia rudis, Madrepora oculata, Paragorgia regalis, Paragorgia* sp. 2, *Paramuricea biscaya, Paramuricea* sp., *Paramuricea* sp. type B3, *Stichopathes* sp. 2, *Swiftia* sp., unknown bamboo 1, and unknown octocoral 1.

Other phylotypes belonging to the SUP05 cluster were present in the water samples and together composed up to 44% of the bacterioplankton in some samples. Seven SUP05 phylotypes were found exclusively in water samples. Similarly, SUP05 phylotypes were found in sediment samples and comprised up to 55% of the microbial community. Five phylotypes were exclusively found in sediment samples.

### Phylogenetic positions of SUP05 phylotypes that associate with corals

Phylogenetic analyses using the full-length and partial 16S rRNA gene sequences revealed that most coral-associated members of the SUP05 cluster belonged to symbiotic Clade 1 as described by Petersen et al (2012) (Fig. 4). This clade includes the chemoautotrophic symbionts of mussels, sponges, snails, and clams from hydrothermal vents, cold seeps, and/or woodfalls. The coral-associated SUP05 in this clade did not form a monophyletic group and were instead placed among many divergent lineages within the clade. Further, most SUP05 phylotypes detected in the water column clustered with free-living members of the SUP05 cluster. Similarly, most SUP05 phylotypes detected in the sediment formed a well-supported cluster (100% UFBoot) near the symbionts of vesicomyid clams.

**Figure 4.**
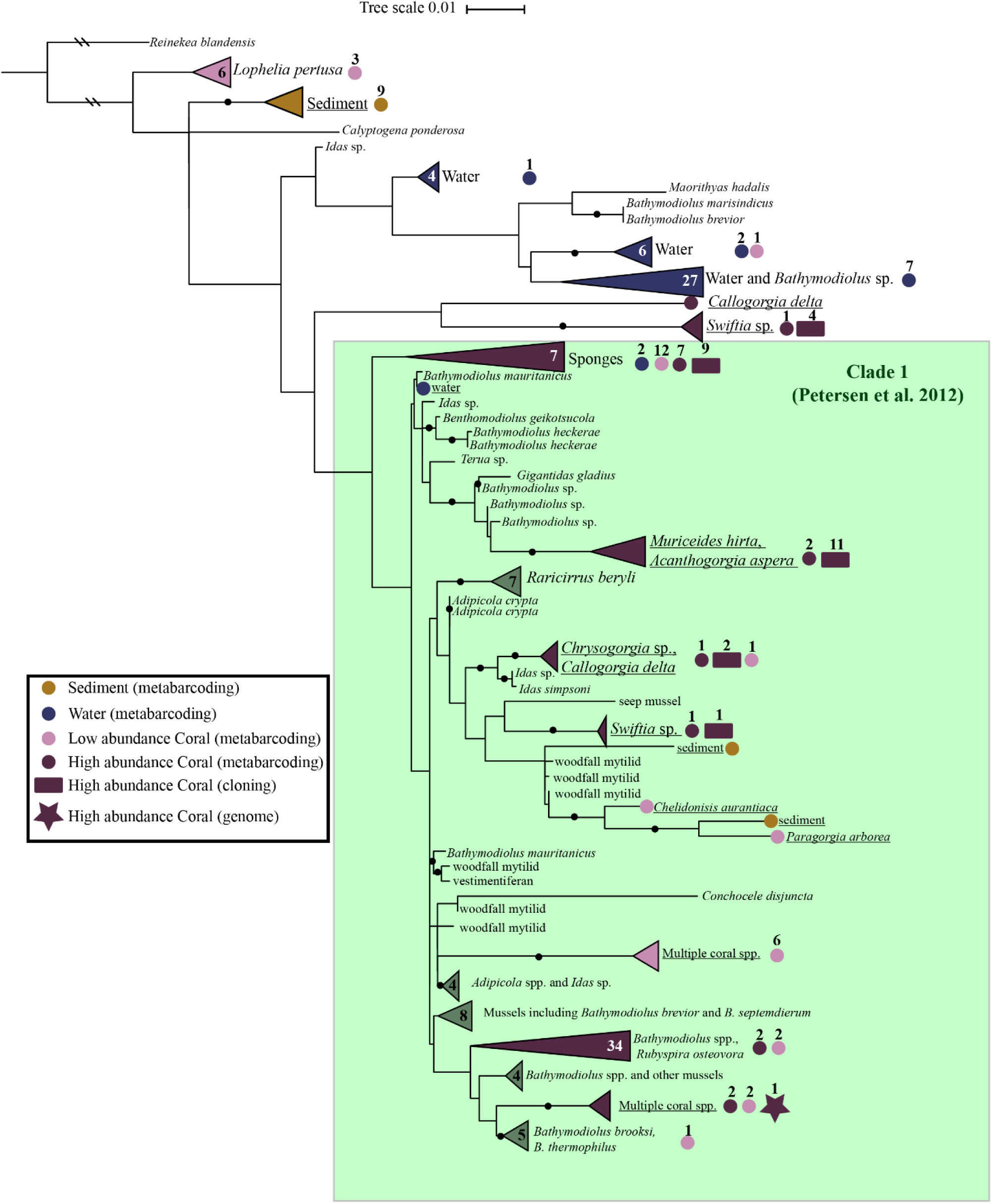
Maximum likelihood phylogenetic tree of all SUP05 16S sequences generated in this study. SUP05 sequences are labeled by their environment: animal host, sediment, or water. Numbers inside collapsed nodes represent the number of database sequences included. Collapsed node labels are underlined if all sequences were generated in this study. Colored circles show the positions of all SUP05 phylotypes based on 16S metabarcoding, rectangles represent the full length 16S sequences generated by cloning, and a star represents the full-length 16S sequences obtained from the genome of the dominant SUP05 from *Paramuricea* sp. type B3. SUP05 sequences colored blue were associated with water samples, brown is associated with sediment samples, and purple is associated with coral samples. High abundance SUP05 phylotypes (>10%) were colored dark purple while low abundance phylotypes were colored light purple.

### Occurrence of SUP05 phylotypes at AT357

A single phylotype of SUP05 was dominant in *Paramuricea* sp. type B3 colonies at AT357 and ranged from 6-85% of the microbial community of the coral (n=22, Fig. 5). This phylotype was also present in three of the four *Madrepora oculata* and one of the three zoanthid gold coral colonies sampled at this site, but only constituted a maximum of 0.069% of their microbial communities. This phylotype was also found in four of the ten sediment samples from this site, with a maximum relative abundance of 0.14% of the sediment community. This phylotype was not found in any water sample from AT357, although other SUP05 phylotypes were present that together comprised between 2.5 - 5.4% of the water microbial community.

**Figure 5.**
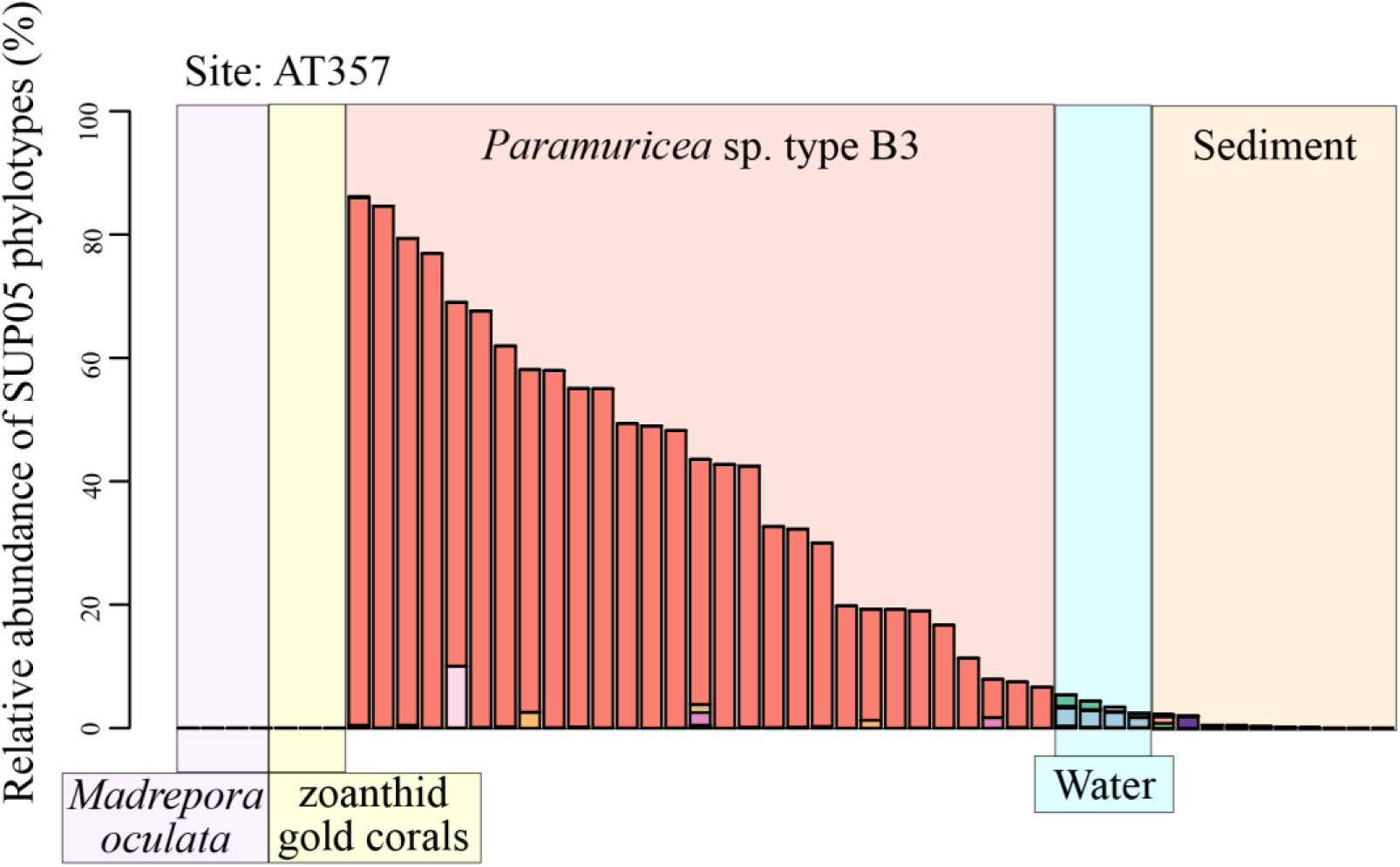
The occurrence of SUP05 phylotypes in corals, water, and sediment at site AT357. The relative abundances of only the phylotypes belonging to the SUP05 cluster are shown. Each color represents a different SUP05 phylotype.

### Metabolic capabilities of a coral-associated SUP05

A 1.9 Mbp draft genome of a member of the SUP05 cluster was assembled from a *Paramuricea* sp. type B3 metagenome. This genome contained the typical gene repertoire for members of the SUP05 cluster and revealed the genetic potential for sulfur-oxidation and carbon fixation. The genes necessary to oxidize the reduced sulfur species thiosulfate, elemental sulfur, and hydrogen sulfide were all present in the genome. These genes included those of the sox pathway (*soxXYZAB*), subunits of dissimilatory sulfite reductase (*dsrA, dsrB*), subunits of adenylyl-sulfate reductase (*apsA, apsB*), sulfate adenylyltransferase (*sat*), rhodanese sulfurtransferase, and sulfide-quinone reductase (*sqr*) (Fig. 6). Transcripts for all of these genes were detected in both RNA libraries with the exception of rhodanese sulfurtransferase which was only detected in one library.

**Figure 6.**
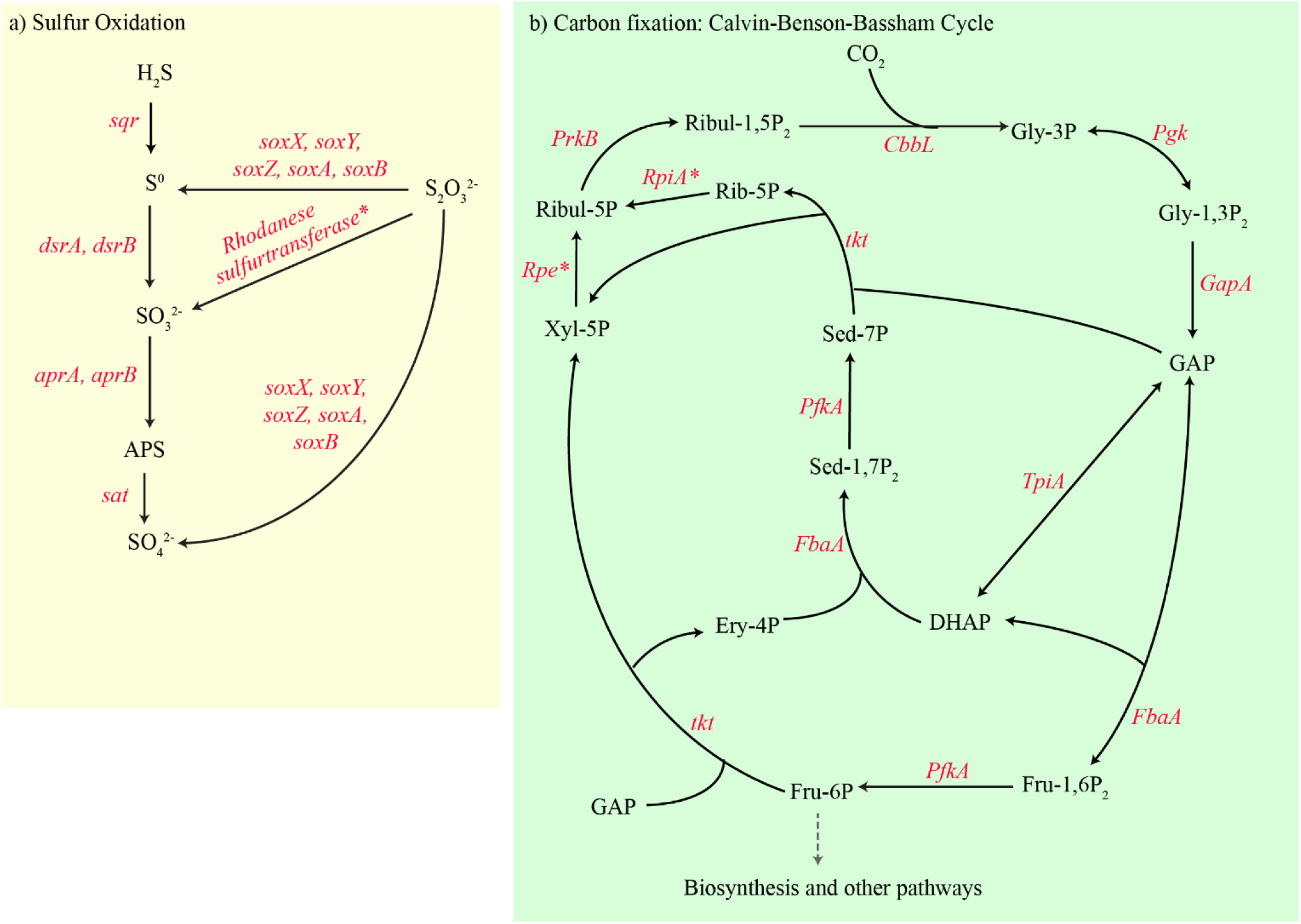
Sulfur oxidation (a) and carbon fixation pathways (b) present in the genome of the dominant SUP05 in *Paramuricea* sp. type B3. Compounds are noted in black text. Genes with transcripts detected in both RNA libraries are denoted in red italics. A * denotes genes with transcripts detected in only 1 RNA library. Abbreviations: APS = adenylyl sulfate; *sqr =* sulfide quinone reductase; *dsrAB* = dissimilatory sulfite reductase; *aprAB* = adenylyl sulfate reductase; *sat* = sulfate adenylyl transferase; *soxXYZAB* = sox pathway genes; Gly-3P = glycerate-3-phosphate; Gly-1,3P_2_ = 1,3-bisphosphoglycerate; DHAP = dihydroxyacetone; GAP = glyceraldehyde phosphate; Fru-1,6P_2_ = fructose-1,6-bisphosphate; Fru-6P = fructose 6-phosphate; Ery-4P = erythrose 4-phosphate; Xyl-5P = xylulose 5-phosphate; Sed-1,7P_2_ = sedoheptulose-1,7-bisphosphate; Sed-7P = sedoheptulose 7-phosphate; Ribul-5P = Ribulose 5-phosphate; Ribul-1,5P_2_ = ribulose-1,5-bisphosphate; Rib-5P = ribose 5-phosphate; *CbbL* = ribulose carboxylase large and small chains; *PgK* = phosphoglycerate kinase; *GapA* = glyceraldehyde-3-phosphate dehydrogenase; *FbaA* = fructose-bisphosphate aldolase; *TpiA* = triosephosphate isomerase; *PfkA* = pyrophosphate-dependent fructose 6-phosphate 1-kinase; *tkt* = transketolase; *Rpe* = Ribulose-phosphate 3-epimerase; *RpiA* = ribose 5-phosphate isomerase A,; *PrkB* = phosphoribulokinase.

The genome also included the genes necessary to fix carbon through a modified Calvin-Benson-Bassham cycle. These genes included the large and small subunits of ribulose bisphosphate carboxylase, phosphoribulokinase, phosphoglycerate kinase, ribulose-phosphate 3-epimerase, glyceraldehyde-3-phosphate dehydrogenase, pyrophosphate-dependent 6-phosphofructokinase, transketolase, fructose-bisphosphate aldolase, triose phosphate isomerase, and ribose-6-phosphate isomerase. The genome was missing sedoheptulose-1,7-bisphosphatase and fructose-1,6-bisphosphatase which are also absent in other SUP05. However, the pyrophosphate-dependent 6-phosphofructokinase encoded in this genome has been suggested to perform the roles of these missing genes in chemoautotrophs (Kleiner et al. 2012; Ponnudurai et al. 2016; Spietz et al. 2019). Transcripts of all of these genes were detected in both RNA libraries except ribose-6-phosphate isomerase and ribulose-phosphate 3-epimerase which were present in only one library each.

Genes involved in other nutritional roles were identified in this SUP05 genome. These genes included assimilatory nitrate and nitrite reductases, genes involved in the biosynthetic pathways of each essential amino acid, and genes with roles in the synthesis of vitamins including biotin, thiamin, folate, and riboflavin.

### Correlation between stable isotopic composition and the relative abundance of SUP05

The tissue stable carbon isotopic compositions of most *Paramuricea* sp. type B3 colonies were within the range of carbon fixed by surface phytoplankton (−22 to −15 ‰ δ ^13^C) however four colonies were more negative with the lowest at −25.6 ‰ (Gearing et al. 1984). The relative abundances of the dominant SUP05 phylotype was negatively correlated with both δ ^13^C and δ ^15^N values (p=0.032, Fig. 7). However, the relative abundance of SUP05 was much more predictive of nitrogen (p=0.00156) than carbon (p=0.08) isotopic compositions.

**Figure 7.**
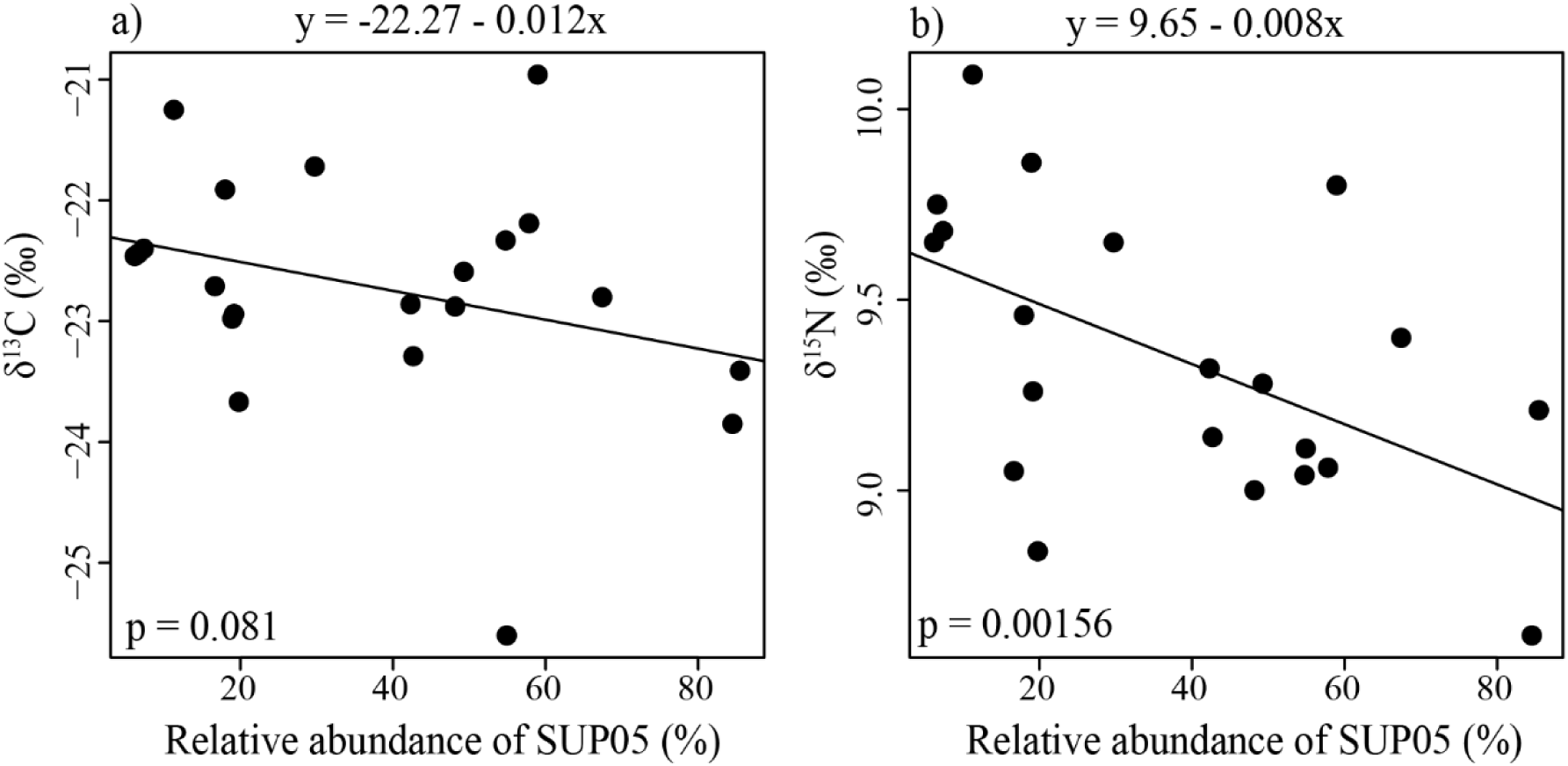
The relationship between the relative abundance of the dominant SUP05 in *Paramuricea* sp. type B3 and the stable carbon (a) and nitrogen (b) isotopic compositions of the coral tissue at site AT357.

## Discussion

### SUP05 is a diverse and abundant taxon that associates with multiple deep-sea coral species

The SUP05 cluster contained a large number of distinct phylotypes and was widespread throughout the deep Gulf of Mexico where we detected it in sediment, water, and multiple species of deep-sea scleractinians, antipatharians and octocorals (Fig. 3). The SUP05 phylotypes most likely to play an important role in corals are those which occur at high relative abundance (>10%). Such abundant SUP05 phylotypes were not found in any shallow or mesophotic corals. Instead they were primarily found in corals that grew at deep sites with signs of active seepage, with the exception of a single coral with 11% SUP05 which was collected near Roberts Reef (Roberts and Kohl 2018), a location without signs of active seepage within lease block VK906. At multiple sites, active signs of seepage were visible within 1 m of some corals. At other sites, no signs of seepage were visible in close proximity to the sampled corals however bacterial mats or seep communities were noted in other unsampled areas of these sites. Thus, coral-associated SUP05 occurred most frequently at sites where corals grew in the vicinity of active signs of seepage.

### Coral-associated SUP05 are probably symbionts

We propose that members of the SUP05 cluster form a symbiotic association with deep-sea corals. First, individual SUP05 phylotypes dominated the microbiome of several coral species and most of those phylotypes were not found in the surrounding sediment or water. This suggests that corals are the main habitat for many SUP05 phylotypes and are not passive contamination from the sediment or water. Additionally, we found several examples of an abundant phylotype of SUP05 that associated with a single species of coral with high fidelity, occurring in this species at multiple sites including sites where other coral-associated phylotypes were present in other coral taxa (Fig. 3).

The location of SUP05 within corals remains uncertain. It seems unlikely that SUP05 inhabits the mucus since most of the abundant phylotypes were not detected in water samples, these octocorals have only a thin layer of mucus, and they were rinsed before preservation. However, microscopy is required to confirm the location of these symbionts.

While SUP05 constituted >10% of the microbiome of many octocorals, *Stichopathes* sp. 3 was the only hexacoral that harbored SUP05 at this level. This may be due to general differences in morphology between octocorals and hexacorals. Most octocorals were erect with smaller polyps and a greater surface area to volume ratio compared to many hexacorals. The morphology of most hexacorals like *Lophelia pertusa* reflect the morphological and physiological restrictions that Childress and Girguis (2011) proposed should exclude an association with sulfur-oxidizing bacteria: thick tissue restricting diffusion of oxygen and lack of physiological adaptations to transport and deliver oxygen.

### The metabolic capabilities of coral-associated SUP05 and its role in corals

We propose that the member of SUP05 that associated with *Paramuricea* sp. type B3 oxidizes reduced sulfur compounds and fixes carbon via the Calvin-Benson-Bassham Cycle like other members of the SUP05 cluster. First, this SUP05 has the genetic potential to conduct these metabolic processes. Second, the genes involved were transcriptionally active. Finally, there were signs of reduced sulfur species in the vicinity of corals that hosted SUP05.

We further hypothesize that *Paramuricea* sp. type B3 near seeps is acquiring some nutrition from the chemoautotrophic primary productivity of its associated SUP05. Colonies with a higher relative abundance of SUP05 also had more chemoautotrophic input into the coral holobiont. Corals may be digesting a small percentage of a resident SUP05 population as *Bathymodiolus* spp. do with their SUP05 symbionts (Fiala-Médioni et al. 1986) or there may be selective transfer of a limited number of organic compounds from the symbiont to host. Although, the stable isotope analysis suggests this nutritional input is quite limited, it may still be very important to the coral host since deep-sea corals are very long-lived (>500 years for some *Paramuricea* spp.) and are food-limited (Girard et al. 2019; Prouty et al. 2016; Roark et al. 2009).

Through nitrate assimilation, this bacterium may be supplying the coral with needed nitrogen because corals mainly eat nitrogen-poor marine snow. This may reflect the stronger correlation between nitrogen and the relative abundance of SUP05 in *Paramuricea* sp. type B3. This SUP05 could also be providing the coral with essential amino acids or vitamins which are not abundant in marine snow. Genes necessary to synthesize all essential amino acids were present as were genes to synthesize four vitamins.

However, although our data strongly suggests a strong relationship between SUP05 and the coral hosts, it does not demonstrate an unequivocal nutritional link. For example, the isotopic signal may simply arise from the bacteria themselves. If SUP05 were very abundant within the coral tissue or mucus but passed no organic compounds to its coral host, then an isotopic signal could arise from the carbon and nitrogen from that the bacteria contribute to each sample. Alternatively, the stable isotope data may reflect nutritional input from other chemoautotrophically-based food sources such as resuspended bacteria from mats or small invertebrates that graze on mats. Finally, SUP05 may simply serve to detoxify hydrogen sulfide in coral tissue by oxidizing it into elemental sulfur.

### Evolutionary history of SUP05 in association with corals

The coral-associated members of the SUP05 cluster did not form a monophyletic group but rather belonged to multiple clades within the cluster. This is similar to the phylogeny of the SUP05 symbionts of Bathymodiolin mussels (Petersen et al. 2012). This suggests that like the association with Bathymodiolin mussels, associations between corals and members of the SUP05 cluster may have evolved multiple times.

### Conclusion

This work has revealed a previously unknown and unexpected association between deep-sea corals and bacteria from the widespread and ecologically important SUP05 cluster. These bacteria exist throughout the world’s oceans where they strongly influence biogeochemical cycles and associate with foundation species at hydrothermal vents and cold seeps. We provide the first indications that this group includes symbionts of deep-sea corals as well which form the foundations of additional diverse animal communities in the deep-sea. This association represents the first association between cnidarians and chemoautotrophic symbionts and may prove important to the ecological functioning of coral communities where they overlap with cold seeps by supplying needed nutrients to food-limited communities.

## Acknowledgements

We would like to thank the crews and pilots of all vessels and ROVs, Dr. Erik Cordes and Dr. Santiago Herrara who served as chief scientists for cruises that made this work possible, and Dr. Peter Etnoyer for providing *Leiopathes glaberrima* samples from the West Florida Slope. We would also like to thank everyone else who helped at sea and in the field: Dr. Fanny Girard, Alexis Weinnig, Alaina Weinheimer, Steve Auscavitch, Dr. Carlos Gomez, Dr. Cherisse Du Preez, Dr. Amanda Glazier, Dr. Jill Bourque, Dr. Amanda Demopoulos, Alicia Yang, Colin Bashaw, Jesse Mee. We would also like to thank Meghann Devlin-Durante for help with extractions and training undergraduates.

## Data Availability

Raw sequence data for metagenomes, the 16S survey, and metatransciptomes were deposited in National Center for Biotechnology Information (NCBI) Sequence Read Archive (SRA) under BioProject numbers PRJNA565265 and PRJNA574146. Data are also publicly available through the Gulf of Mexico Research Initiative Information & Data Cooperative (GRIIDC) at https://data.gulfresearchinitiative.org (doi:<10.7266/RYMQTDQ9>, R4.x268.000:0125, and R4.x268.000:0125)

